# K-space sampling strategies to reduce noise induced by cardiac pulsatility in brain maps of R_2_^*^ and magnetic susceptibility

**DOI:** 10.1101/2024.08.20.608759

**Authors:** Quentin Raynaud, Thomas Dardano, Rita Oliveira, Giulia Di Domenicantonio, Tobias Kober, Christopher W. Roy, Ruud B. van Heeswijk, Antoine Lutti

## Abstract

Maps of the transverse relaxation rate R2* and magnetic susceptibility (𝜒) are computed from gradient-echo data acquired at multiple echo times and are sensitive to signal instabilities induced by cardiac pulsation. Here, we introduce two k-space sampling strategies that aim to mitigate the impact of cardiac-induced noise in brain maps of R2* and 𝜒.

The proposed strategies are based on the higher level of cardiac-induced noise near the k-space centre compared to the periphery. Using a CArtesian trajectory with Spiral PRofile (CASPR), the first strategy allows for the acquisition of a specific number of averages at each k-space location, derived from the local level of cardiac-induced noise. The second strategy synchronizes the acquisition near the k-space centre with the cardiac cycle in real time. We compared the variability across 4 repetitions of R2* and 𝜒 maps computed from data acquired using both strategies and with a standard linear trajectory.

Data was acquired in 10 healthy volunteers. Compared to linear trajectory, the CASPR trajectory reduced the variability of R2* and 𝜒 maps across repetitions by 26/28/22% and 19/18/16% in the brainstem/cerebellum/whole brain, for a 14% increase in scan time. The CASPR trajectory also reduced the level of aliasing artifacts from pulsating blood vessels. The synchronized trajectory did not reduce the variability of R2* or 𝜒 maps.

CASPR trajectories can be designed to mitigate cardiac-induced noise in brain maps of the MRI parameters R2* and 𝜒. Synchronization of data acquisition with the cardiac cycle did not reduce the level of cardiac-induced noise.

## 1. Introduction

MRI relaxometry enables the non-invasive investigation of microscopic changes in brain tissue in patient populations.^1–3^ Estimates of the transverse relaxation rate R2* (=1/T2*)^4^ and magnetic susceptibility (𝜒)^5^ are biomarkers of iron and myelin concentration within brain tissue,^3,6,7^ and provide means to monitor the evolution of neurological diseases such as Parkinson’s disease,^8,9^ Alzheimer’s disease,^10^ and multiple sclerosis^11^ in patient populations.

R2* and 𝜒 measurements can both be perturbed by cardiac-induced noise. Cardiac pulsation leads to a systolic blood pressure wave that travels to the brain and generates periodic signal instabilities through head motion,^12^ brain tissue deformation,^13,14^ blood flow,^15–18^ cerebrospinal fluid flow,^19,20^ changes in the tissue’s O2/CO2 concentration,^21,22^ blood vessel pulsatile motion^23^ and other effects.^24^ Cardiac pulsation leads to signal instabilities that are most prominent in inferior and highly vascularized brain regions such as the orbitofrontal cortex,^20,25^ brainstem,^26,27^ cerebellum^28^ and periventricular regions.^25,29^ Changes of the B0-field,^30^ laminar flow^31,32^ and motion^33,34^ are coherent across an image voxel and lead to a net phase shift of the signal. Turbulent flow^35^ and intra-voxel B0 inhomogeneities^36^ are incoherent across an image voxel and affect the magnitude of the signal. Noise induced by cardiac pulsation increases with the echo time,^37,38^ which commonly reaches ∼40ms in the gradient-recalled echo (GRE) data used for the computation of R2* and 𝜒 maps.^11,39,40^ As a result, cardiac-induced noise increases the variability of R2* and 𝜒 maps across repetitions, reducing sensitivity to brain change in neuroscience studies.^40,41^ Cardiac-induced noise also leads to aliasing artifacts in inferior or highly vascularized brain regions.^42^

In functional MRI, series of image volumes are acquired at a rate of ∼1Hz and the effect of cardiac pulsation on the time course of the data can be removed using dedicated models.^43–45^ In MRI relaxometry, data acquisition requires several minutes and the effects of cardiac-induced signal instabilities are challenging to reduce with post-processing techniques. Instead, prospective strategies have been proposed to ensure that cardiac-induced noise remains incoherent along the spatial^46–48^ or temporal^42^ dimensions of the multi-echo GRE data. These passive scrambling methods, which aim to mitigate the effect of cardiac-induced noise rather than reduce its amplitude, remove aliasing in R2* and 𝜒 maps and improve their reproducibility by ∼25%.^42^ Strategies that reduce the level of cardiac-induced noise in GRE data target in-flowing arterial blood.^38,49^ However, these techniques only address a subset of the physiological effects that contribute to cardiac-induce noise. They also require tailored radiofrequency (RF) saturation pulses or gradient waveforms that reduce the efficiency of data acquisition. Instead, reduction of cardiac-induced noise in GRE data may be achieved from alternative k-space sampling trajectories: keyhole imaging or PROPELLER acquire the centre of k-space multiple times to resolve dynamic effects such as blood flow^50^ or BOLD signals^51^ and are tailored to increase SNR or reduce motion.^52,53^ In non-brain imaging, dynamic keyhole acquisition can be used to resolve and mitigate breathing effects or motion.^54–56^ The adaptation of these methods towards the reduction of cardiac-induced noise in GRE data remains unexplored.

In this study, we introduce two data acquisition strategies that aim to reduce the level of cardiac-induced noise in brain relaxometry data. These strategies were derived from a recent characterization of cardiac-induced noise in multi-echo GRE data.^41^ The first strategy (CArtesian trajectory with Spiral PRofile ordering, CASPR),^56^ makes use of the acquisition of a specific number of averages at each k-space location, set according to the local level of cardiac-induced noise. The second strategy, inspired by the use of cardiac gating for diffusion^57,58^ and functional MRI,^59,60^ synchronizes data acquisition near the k-space centre with cardiac pulsation to acquire the most sensitive data during the most stable part of the cardiac cycle. Following the optimization of both strategies using numerical simulations, we evaluated their ability to reduce the level of cardiac-induced noise in repeated multi-echo GRE data acquired *in vivo*.

## 2. Methods

The proposed strategies to reduce the level of cardiac-induced noise were derived from a recent characterization of cardiac-induced noise in GRE data.^41^ In that study, low-resolution 3D multi-echo GRE data was acquired continuously for one hour in 5 participants. The participants’ cardiac pulsation was recorded simultaneously. To resolve the effect of cardiac pulsation on R2* maps, the data was reordered according to the phase of the cardiac cycle at the time of its acquisition, leading to 5D datasets with three spatial dimensions, one dimension for the echo time and one dimension for the phase of the cardiac cycle. By analogy with other models of physiological noise,^43,45,61^ cardiac-induced noise was modelled by second-order Fourier series decomposition of the change of the 5D data along its cardiac cycle dimension.^41^ This enabled the estimation of the amplitude of cardiac-induced noise at each of k-space location and echo time.

Here, we used the cardiac-induced noise modelled from the 5D datasets to conduct numerical simulations of candidate k-space sampling strategies and to evaluate their ability to reduce cardiac-induced noise in multi-echo GRE data.

### 2.1. Strategy 1: CArtesian trajectory with Spiral PRofile (CASPR)

The amplitude of the modelled cardiac-induced noise depends primarily on the radial distance |𝑘| to the centre of k-space along the two phase-encoding directions (𝑘_𝑦_, 𝑘_𝑧_) of a 3D GRE sequence (Figure 1A): 50-60% of cardiac-induced noise is located near the k-space centre (|𝑘| < 0.074 mm^−1^).^41^ With Strategy 1, the number of averages at a distance |𝑘| from the k-space centre is set according to the local level of cardiac-induced noise. The effective noise level in the data is:

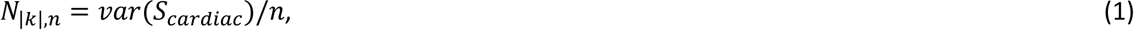

**Figure 1:**
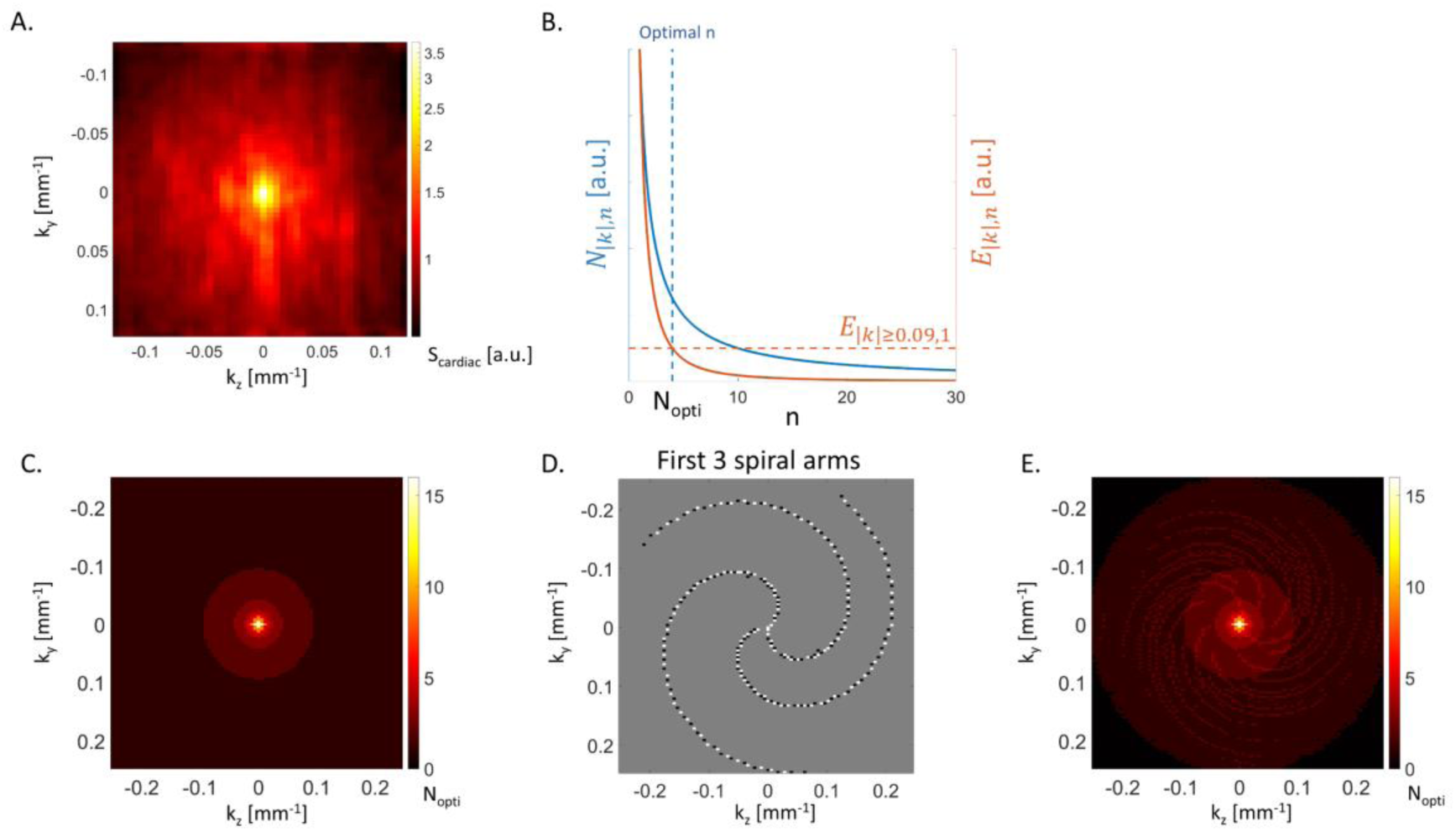
Proposed CASPR trajectory. (A) Distribution of the amplitude of the modelled cardiac- induced noise 𝑆𝑐𝑎𝑟𝑑𝑖𝑎𝑐 along the two phase-encoding directions (𝑘𝑦, 𝑘𝑧) of a 3D GRE sequence.50 (B) Effective noise level (𝑁|𝑘|,𝑛) and averaging efficiency (𝐸|𝑘|,𝑛) as a function of the number of averages n. The optimal number of averages Nopti at a given k-space location is reached when the averaging efficiency (𝐸|𝑘|,𝑛) equals that of a point at the k-space periphery dominated by thermal noise. (C) Target k-space distribution of Nopti. (D) First 3 spiral arms of the optimal CASPR trajectory. The white/black data points are acquired on the Cartesian grid in the outward/inward direction. (E) Distribution of Nopti achieved with the optimal CASPR trajectory. The k-space corners (in black) are acquired linearly at the end of the CASPR acquisition.

where 𝑛 is the number of averages and 𝑣𝑎𝑟(𝑆_𝑐𝑎𝑟𝑑𝑖𝑎𝑐_) is the variance of the modelled cardiac-induced signal fluctuations across the cardiac cycle^41^.

The averaging efficiency, i.e. the reduction in effective noise level from the acquisition of one additional average, is given by:

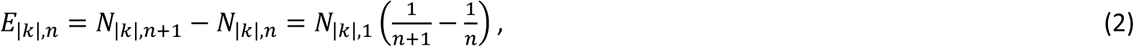

where 𝑁_|𝑘|,1_ = 𝑣𝑎𝑟(𝑆_𝑐𝑎𝑟𝑑𝑖𝑎𝑐_) is the effective noise level when only one sample is acquired. Equation 2 shows that averaging efficiency decreases with an increasing number of averages (Figure 1B). Enforcing a uniform effective noise level across k-space (equation 1) would require a high number of averages near the k-space centre. Because efficiency decreases with the number of averages, this would lead to a prohibitive extension of scan time. Instead, we set the optimal number of averages 𝑁_𝑜𝑝𝑡𝑖_ so that the averaging efficiency is at least equal to that of the k-space periphery (|𝑘| ≥ 0.09 mm^-1^), where thermal noise is the dominant source of noise and only one sample is acquired (𝑛 = 1):

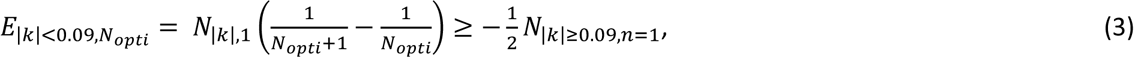

Figure 1C shows the distribution of 𝑁_𝑜𝑝𝑡𝑖_ in the 2D plane of the two phase-encoding directions (𝑘_𝑦_, 𝑘_𝑧_). This distribution, which peaks at 𝑁_𝑜𝑝𝑡𝑖_∼15 near the k-space centre, was achieved using a CArtesian trajectory with Spiral PRofile (CASPR).^56,62^ K-space sampling was conducted on a Cartesian grid, travelling outward and inward along the spiral arms (Figure 1D).^63,64^ CASPR trajectories with different number of points per arms (10 to 200), number of arms (10 to 200) and sampling density (0.1 to 1.75) were simulated. The CASPR trajectory that led to the closest match with the target number of averages consisted of 120 spiral arms with 100 points each (Figure 1E). The spatial frequencies in the corners of k-space (black regions of Figure 1E), not acquired with the CASPR trajectory, are acquired using a linear scheme at the end of the acquisition.

While CASPR is also used with non-Cartesian trajectories,^63–66^ the proposed implementation samples k-space data on a 3D Cartesian grid: the (𝑘_𝑦_, 𝑘_𝑧_) positions are defined on a Cartesian grid and sampling along the readout direction is conducted using a Cartesian linear trajectory, using trapezoidal gradient waveforms. As a result, no data interpolation or non-uniform Fourier transformation is required prior to image reconstruction.

### 2.2. Strategy 2: Cardiac-triggered sampling

Strategy 2 proposes to use cardiac triggering to enforce data acquisition during the part of the cardiac cycle where the GRE signal is most stable. Here, cardiac triggering locks the data acquisition to a fixed phase of the cardiac cycle at specific k-space locations only: preceding and subsequent peripheral k-space locations are sampled without locking. To limit the resulting extension of scan time, cardiac triggering was only applied near the k-space centre (|k| < 0.125mm^-1^), where most cardiac-induced noise is present.^41^

We conducted numerical simulations to identify an implementation of cardiac triggering that leads to maximal reduction of cardiac-induced noise in the acquired data. From the 5D model of cardiac-induced noise, we computed the first order derivative along the direction of the cardiac cycle, then averaged it along the readout (𝑘_𝑥_) and echo time directions, resulting in a 3D matrix containing the derivatives of the modelled cardiac-induced noise along the direction of the cardiac cycle, for each (𝑘_𝑦_, 𝑘_𝑧_) coordinate of the phase-encoding plane. This matrix was used as a fingerprint of signal instability induced by cardiac pulsation, for each (𝑘_𝑦_, 𝑘_𝑧_) coordinate and phase of the cardiac cycle.

The global signal instability is lower during the first quarter of the cardiac cycle (0 < 𝜑_𝑐_ < _2_) (Figure 2A).

**Figure 2:**
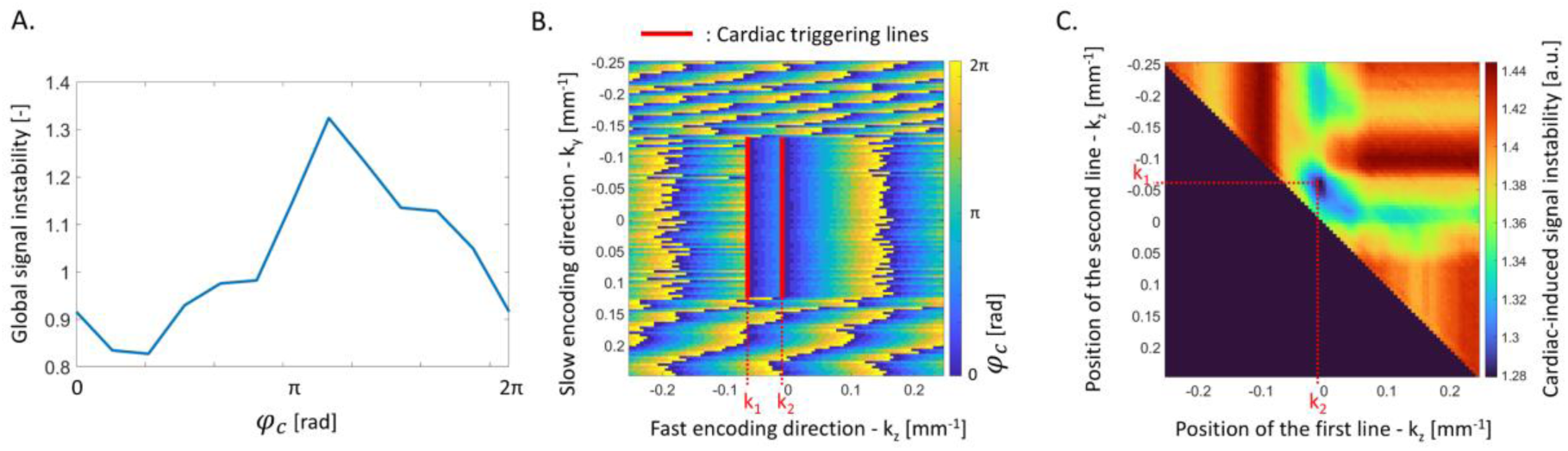
Proposed cardiac-triggered sampling. (A) Global signal instability induced by cardiac pulsation. (B) Phase of the cardiac cycle at the time of acquisition of the data with the proposed cardiac-triggered linear trajectory. The fast (𝑘𝑧)/slow (𝑘𝑦) phase-encoding directions of the 3D GRE sequences are displayed horizontally/vertically. K-space sampling was started from the top-left corner. The red cardiac triggering lines enforce acquisition of the k-space centre data during the most stable part of the cardiac cycle (0 < 𝜑𝑐 < 2). (C) Cardiac-induced noise level as a function of the position of the two cardiac triggering lines 𝑘1 and 𝑘2.

Simulation of data acquisition consisted of selecting one value of cardiac-induced signal instability for each (𝑘_𝑦_, 𝑘_𝑧_) coordinate, depending on the sequence timing and the cardiac period 𝑇, i.e. the time required to span the 2𝜋 range of the cardiac phase dimension of the matrix of cardiac-induced signal instability. The simulated MRI acquisition had a repetition time (TR) of 40ms and a matrix size of 88 along the fast phase-encoding direction (𝑘_𝑧_). K-space sampling was started from one corner of k-space, as a standard linear trajectory, with no locking to the cardiac cycle (Figure 2B). For each consecutive k-space coordinate in the slow phase-encoding direction (𝑘_𝑦_) data was acquired linearly along the fast- phase-encoding direction (𝑘_𝑧_), at a rate of 1/TR = 25Hz. As a result, the phase of the cardiac cycle at the time of acquisition of the data varied smoothly along 𝑘_𝑧_. When the k-space centre was reached (|k| < 0.125mm^-1^), data acquisition was locked to the cardiac cycle by means of two cardiac triggering lines aligned along the slow phase-encoding direction 𝑘_𝑦_ (Figure 2B). Upon reaching these lines, data acquisition was paused, maintaining RF excitation to preserve the steady state of the MRI signal.^67^ K- space sampling was resumed when the next peak of the pulse wave was detected (𝜑_𝑐_ = 0). As a result, acquisition of the subsequent data points took place during the most stable period of the cardiac cycle (i.e. the first quarter after systole). A global estimate of signal instability was computed by summing the selected cardiac-induced signal instabilities across all (𝑘_𝑦_, 𝑘_𝑧_) coordinates. The simulations were repeated, varying the cardiac period 𝑇 from 600ms to 1000ms in steps of 10ms.

The numerical simulations were conducted for every possible position of the two cardiac triggering lines. The optimal position of the cardiac triggering lines, identified from the minimum global signal instability induced by cardiac pulsation, was 𝑘_1_ = −0.0625 mm^-1^ and 𝑘_2_ = −0.0057 mm^-1^ along the fast-phase-encoding direction (Figure 2C).

### 2.3. MRI data acquisition and analyses

To assess the reduction in cardiac-induced noise from the two proposed strategies, data was acquired in 10 healthy volunteers (7 females, 27±7 years old), with a 3T MRI scanner (Magnetom Prisma, Siemens Healthineers, Forchheim, Germany) and a 64-channel head-neck coil. The participants’ cardiac pulsation was recorded using a pulse-oximeter attached to their finger. To ensure optimal quality of the pulse-oximeter signal, participants were offered a blanket and a hot water bottle was placed near their hand. The study was approved by the local ethics committee (CER-VD) and all participants gave their written informed consent prior to participation.

#### 2.3.1. MRI protocol

MRI data were acquired with a custom-made 3D multi-echo FLASH sequence capable of k-space sampling with a standard linear Cartesian trajectory as well as the proposed CASPR trajectory and cardiac triggering. 15 echo images were acquired after radiofrequency (RF) excitation with echo times TE=2.34ms to 35.10ms. The repetition time was 40ms, the RF excitation flip angle was 16° and image resolution was 2x2x2mm^3^ (matrix size 128x128x88). GRAPPA^68^ was used with acceleration factor of 2 and 24 reference lines. The scan time of each repetition was 4:31 minutes for the standard linear trajectory, 5:09 minutes (+14%) for the CASPR trajectory and 5:22 minutes (+19±1%) for the cardiac-triggered sampling. Four repetitions were conducted for each sampling strategy to quantify the reproducibility of the data. The spatial resolution of the data was low to keep the total scan time reasonable. Additional MRI data were acquired on a single participant (male, 30 years old) with a higher image resolution of 1.2x1.2x1.2mm^3^ (matrix size 208x192x144), representative of relaxometry protocols used in neuroscience studies.^69–71^ The other acquisition parameters were unchanged. Three repetitions of the high-resolution protocol were conducted with standard linear trajectory (scan time 10:28 minutes) and CASPR trajectory (scan time 13:30 minutes (+29%)).

All acquisition protocols also included an MP-RAGE^72^ image for segmentation and anatomical reference (1mm^3^ resolution, TR/TE = 2000/2.39ms, GRAPPA^68^ acceleration factor 2 with 24 reference lines, RF excitation angle = 9°, acquisition time 4:16 minutes). Two 3D FLASH datasets were acquired with the head- and body coils for signal reception (4×4×4mm^3^ image resolution, TR/TE=5.72ms/2.34ms, excitation flip angle=6°, acquisition time 16s) and used subsequently for the computation of the coil sensitivity maps.^73^

#### 2.3.2. Image reconstruction

Brain images were reconstructed offline using Matlab (version 2022a, The MathWorks, Natick, MA). Coil sensitivity maps were computed as the ratio of the (4mm)^3^ resolution data acquired with the head and body coils.^73,74^ For CASPR trajectory, the multiple samples acquired at each k-space location were averaged.^75^ Coil-specific images were reconstructed with GRAPPA^68^ (https://github.com/mchiew/grappa-tools) and combined by performing a SENSE^74^ reconstruction with an acceleration factor of 1.

### 2.4. Computation and analysis of the qMRI maps

#### 2.4.1. Relaxometry and QSM

R2* maps were computed voxel-wise from multi-echo images on each repetition using a regression of the log signal with the corresponding echo times.^76^ The noise level on the R2* estimates was calculated as the root-mean-squared error (RMSE) between the MR signal and the R2* fit.

Maps of magnetic susceptibility (𝜒) were generated from the phase of the MR data of each repetition using bespoke scripts adapted from https://github.com/fil-physics/MPM_QSM and following the ISMRM consensus guidelines.^77^ The phase maps were unwrapped using ROMEO^78^ with an additional correction for linear phase offsets induced by bipolar readouts.^79^ Brain masks were generated with BET from FSL^80^ and underwent subsequent refinement through multiplication with a phase-quality-based mask obtained during the ROMEO unwrapping process, and any holes present within the resulting mask were subsequently filled. Removal of the background field was conducted with the Projection onto Dipole Fields algorithm^81^ available in the SEPIA toolbox.^82^ Finally, dipole inversion was computed using the STAR-QSM algorithm^83^ provided in the SEPIA toolbox, using the entire brain as a reference. In one subject, an interhemispheric calcification was manually masked out and dipole inversion was conducted using this adjusted mask.

#### 2.4.2. Image segmentation and statistical analysis

Image coregistration and segmentation were conducted using Statistical Parametric Mapping (SPM12, Wellcome Centre for Human Neuroimaging, London, UK). The MP-RAGE images were segmented into maps of grey and white matter probabilities using Unified Segmentation.^84^ Whole-brain masks were computed from the grey and white matter segments and included voxels with a combined probability of 0.9 or above. As described in Lutti et al.,^69^ regional masks were computed from the grey matter maximum probability labels computed in the ‘MICCAI 2012 Grand Challenge and Workshop on Multi-Atlas Labeling’ (https://masi.vuse.vanderbilt.edu/workshop2012/index.php/Challenge_Details), using MRI scans from the OASIS project (http://www.oasis-brains.org/) and labelled data provided by Neuromorphometrics, Inc. (http://neuromorphometrics.com/) under academic subscription.

Cardiac pulsation leads to exponential-like and non-exponential effects on the magnitude of the MRI signal.^41^ The level of cardiac-induced noise in the magnitude of the MRI data was therefore assessed from the estimates of R2* and RMSE. The level of cardiac-induced noise in the phase of the MRI was assessed from the 𝜒 estimates alone: the noise level on these estimates could not be computed from the code used for the generation of the 𝜒 maps.

Because cardiac-induced noise increases the variability of R2*, RMSE and 𝜒 estimates,^41,42^ the standard deviation (SD) of these metrics across repetitions was used in the statistical comparisons of the proposed acquisition strategies. Additionally, an increase of the mean RMSE across repetitions has been reported as an indicator of the successful mitigation of the exponential-like effects of cardiac pulsation on the signal magnitude.^42^ Statistical comparisons of the proposed acquisition strategies therefore also included the mean RMSE. These comparisons were conducted using two-sided Student’s t-tests of regional averages computed from the brainstem and cerebellum – inferior brain regions that exhibit large cardiac-induced effects - and the whole-brain.

## 3. Results

With the standard linear trajectory, the SD of R2* across repetitions reaches up to 4s^-1^ in inferior brain regions (Figure 3A). With the CASPR trajectory, maps of the SD of R2* are more spatially uniform and the regional averages are reduced by 26/28/22% in the brainstem/cerebellum/whole-brain compared to the standard linear trajectory (Table 1 and Figure 3, p≤0.017). This is larger than the √1.14 ∼ 6.7% reduction in variability expected from the 14% additional samples of the CASPR trajectory if thermal noise dominated signal variance. With cardiac triggering, the SD of R2* across repetitions is reduced by 6/6/3% in the brainstem/cerebellum/whole-brain respectively, for an increase in scan time of ∼19% (Figure 3, p≥0.52). This is less than the reduction in variability expected from data acquired with the standard linear trajectory with 19% more samples to improve SNR (√1.19 ∼ 9.0%). Post-hoc analyses were conducted to verify that the underwhelming reproducibility of the data acquired with cardiac triggering is not due to a faulty synchronization of data acquisition with the participants’ cardiac cycle (see Supplementary Material S1).

**Figure 3:**
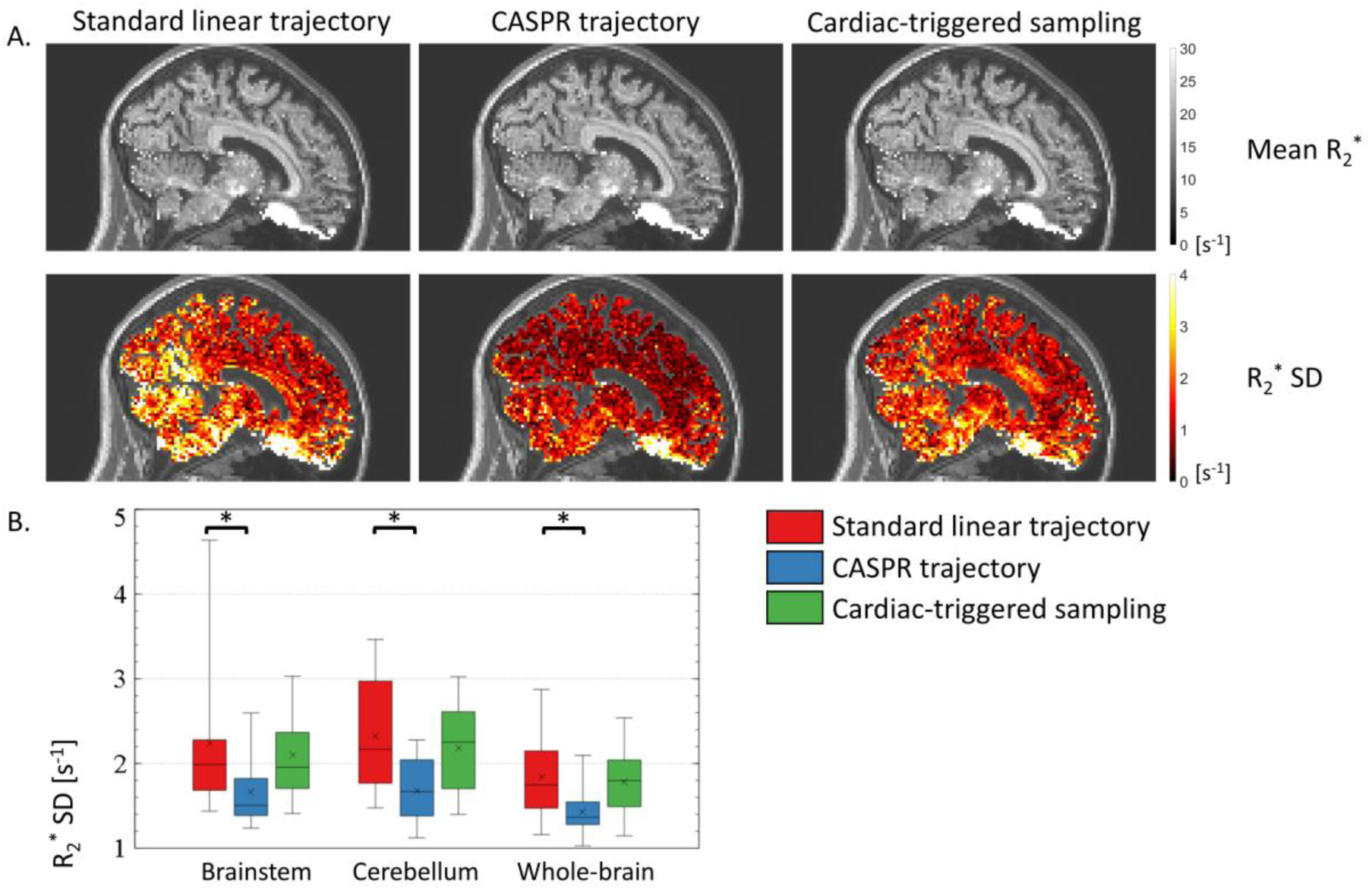
(A) Example maps of the mean and standard deviation (SD) of R2* across repetitions, computed from data acquired with the standard linear and CASPR trajectories, and with cardiac triggering. (B) Regional estimates of the SD of R2* across repetitions in the brainstem, cerebellum and whole brain, across all 10 healthy volunteers.

**Table 1:** ROI-averages of the variability of the R2*/RMSE/𝜒 and mean RMSE estimates across repetitions, computed from data acquired with the standard linear and CASPR trajectories and with cardiac triggering.

|  | Sampling | Brainstem | Cerebellum | Whole brain |
| --- | --- | --- | --- | --- |
| $R_2^*$ SD [$s^{-1}$] | Standard linear | 2.24±0.93 | 2.33±0.73 | 1.84±0.52 |
|  | CASPR | 1.67±0.43 | 1.68±0.41 | 1.43±0.32 |
|  | Cardiac triggering | 2.10±0.55 | 2.18±0.56 | 1.79±0.43 |
| RMSE mean [a.u.] | Standard linear | 3.87±0.78 | 3.81±0.71 | 2.90±0.39 |
|  | CASPR | 3.69±0.58 | 3.70±0.54 | 2.84±0.33 |
|  | Cardiac triggering | 3.92±0.46 | 3.88±0.53 | 2.95±0.28 |
| RMSE SD [a.u.] | Standard linear | 0.98±0.32 | 0.95±0.26 | 0.69±0.14 |
|  | CASPR | 0.82±0.13 | 0.79±0.15 | 0.60±0.09 |
|  | Cardiac triggering | 0.94±0.16 | 0.95±0.21 | 0.69±0.12 |
| $\chi$ SD [ppm] $\times 10^3$ | Standard linear | 9.03±3.27 | 9.74±2.79 | 7.66±2.01 |
|  | CASPR | 7.41±2.68 | 8.02±2.10 | 6.50±1.56 |
|  | Cardiac triggering | 8.66±2.41 | 9.26±1.76 | 7.33±1.30 |

The maps of the mean and SD of RMSE across repetitions appear largely comparable between the three candidate sampling strategies (Figure 4A). Compared to standard linear trajectory, the mean RMSE is lower by 5/3/2% with the CASPR trajectory (p≥0.06) and higher by 1/2/1% with cardiac triggering (p≥0.33), in the brainstem/cerebellum/whole-brain, respectively (see Table 1 and Figure 4B). With the CASPR trajectory, the SD of RMSE across repetitions is lower by 17/17/13% in the brainstem/cerebellum/whole-brain compared to standard linear trajectory (Figure 4B, p≤0.046). With cardiac triggering, the SD of RMSE across repetitions is higher by 5/1/0% (Figure 4B, p≥0.58).

**Figure 4:**
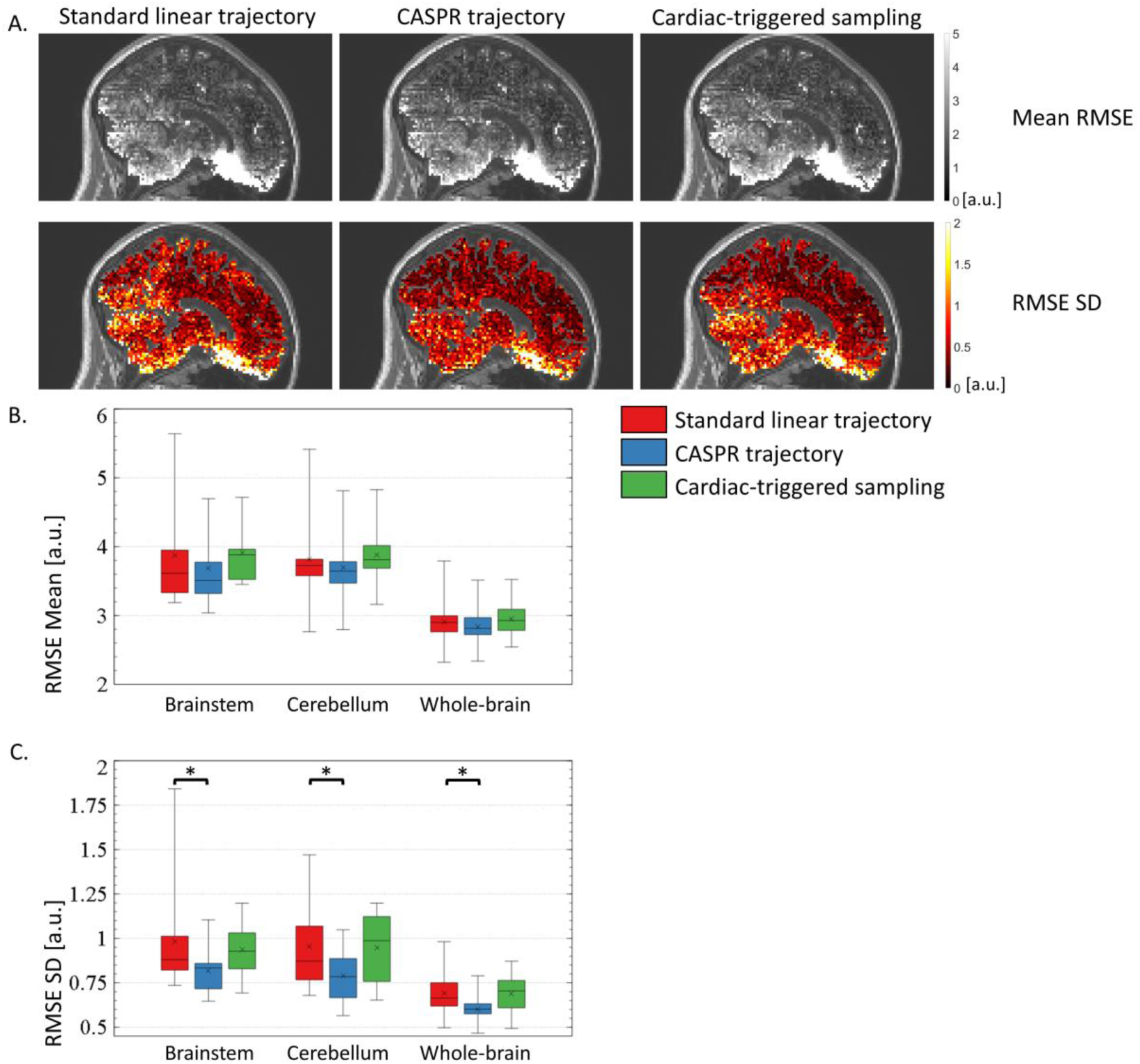
(A) Example maps of the mean and standard deviation (SD) of RMSE across repetitions, computed from data acquired with the standard linear and CASPR trajectories, and with cardiac triggering. (B) Regional estimates of the mean RMSE in the brainstem, cerebellum and whole brain, across all 10 healthy volunteers. (C) Regional estimates of the SD of RMSE in the brainstem, cerebellum and whole brain, across all 10 healthy volunteers.

The maps of the mean and SD of 𝜒 across repetitions display the same spatial features between the three candidate sampling strategies (Figure 5A). With the CASPR trajectory, the SD of the 𝜒 estimates across repetitions is lower than that of the standard linear trajectory by 19/18/16% in the brainstem/cerebellum/whole-brain (Figure 5B, p≤0.015). With the cardiac triggering, the SD of the 𝜒 estimates across repetitions is lower by 4/5/4% (Figure 5B, p≥0.38).

**Figure 5:**
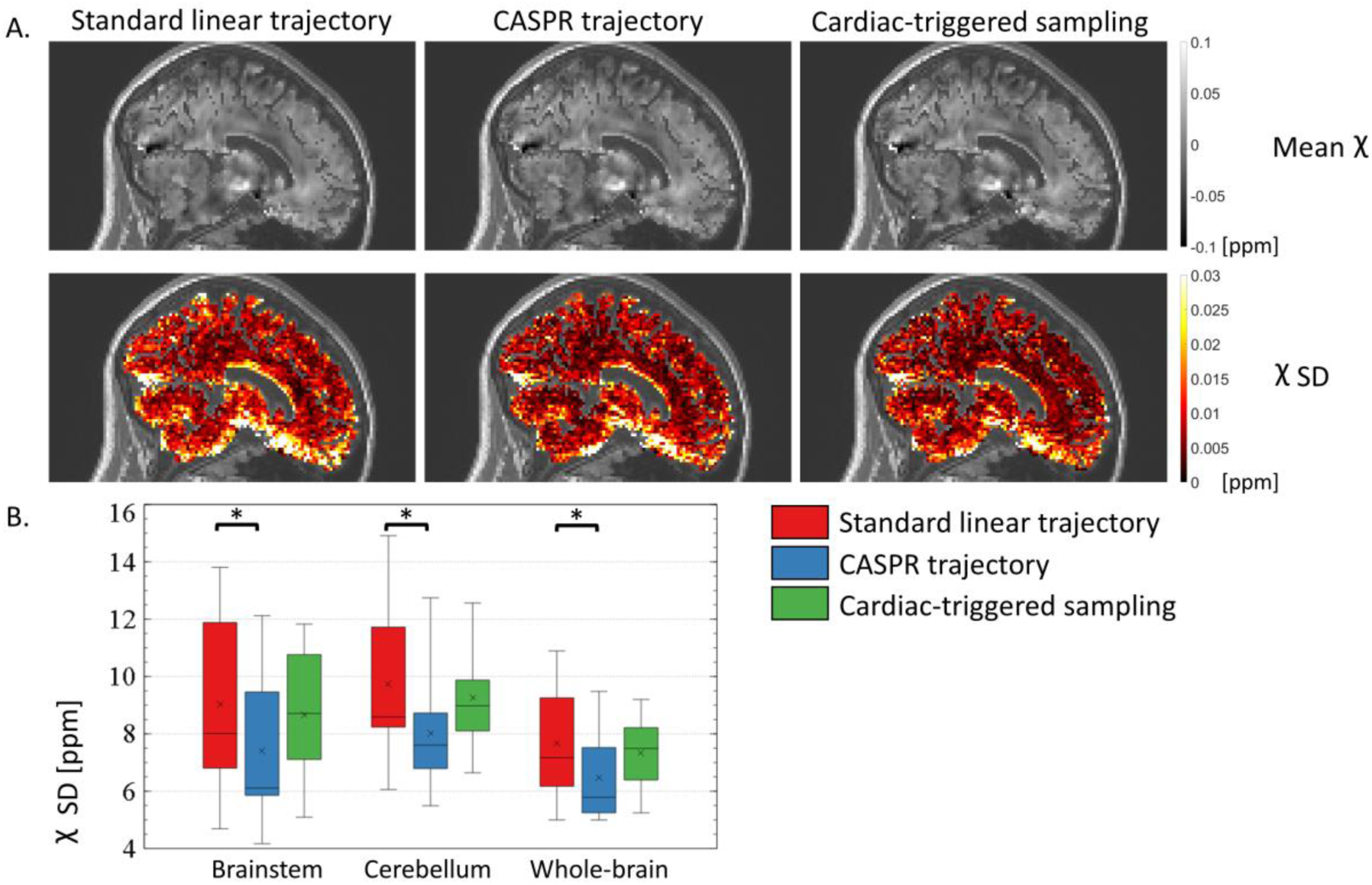
(A) Example maps of the mean and standard deviation (SD) of 𝜒 across repetitions, computed from data acquired with the standard linear and CASPR trajectories, and with cardiac triggering. (B) Regional estimates of the SD of 𝜒 across repetitions in the brainstem, cerebellum and whole brain, across all 10 healthy volunteers.

The high-resolution data acquired with the standard linear trajectory shows a clear pattern of increased variability across repetitions in inferior brain regions (Figure 6A). In particular, maps of the SD of R2* show aliasing along the anterior-posterior direction of cardiac-induced noise originally located e.g., in the Circle of Willis. This aliasing is not present in the data acquired with the CASPR trajectory. The SD of R2* across repetitions is lower by 31/36/36% in the brainstem/cerebellum/whole-brain with the CASPR trajectory than the standard linear trajectory. The SD of the 𝜒 estimates is lower by 27/26/32% with the CASPR trajectory than the standard linear trajectory in the brainstem/cerebellum/whole-brain (Figure 6B). The reduction of the SD of R2* and 𝜒 maps across repetitions is larger than the √1.29 ∼ 13.6% reduction expected from the 29% of additional samples of CASPR trajectory if thermal noise dominated signal variance. R2* maps computed from data acquired with the standard linear trajectory also show aliasing of cardiac-induced noise along the anterior-posterior direction (Figure 6C, blue arrows). With the CASPR trajectory, this aliasing is not present, and the delineation of the white-grey matter is improved. The 𝜒 data acquired with linear trajectory show large streaking artifacts^83^ around the Circle of Willis that are strongly reduced with CASPR trajectory (Figure 6D, green arrows).

**Figure 6:**
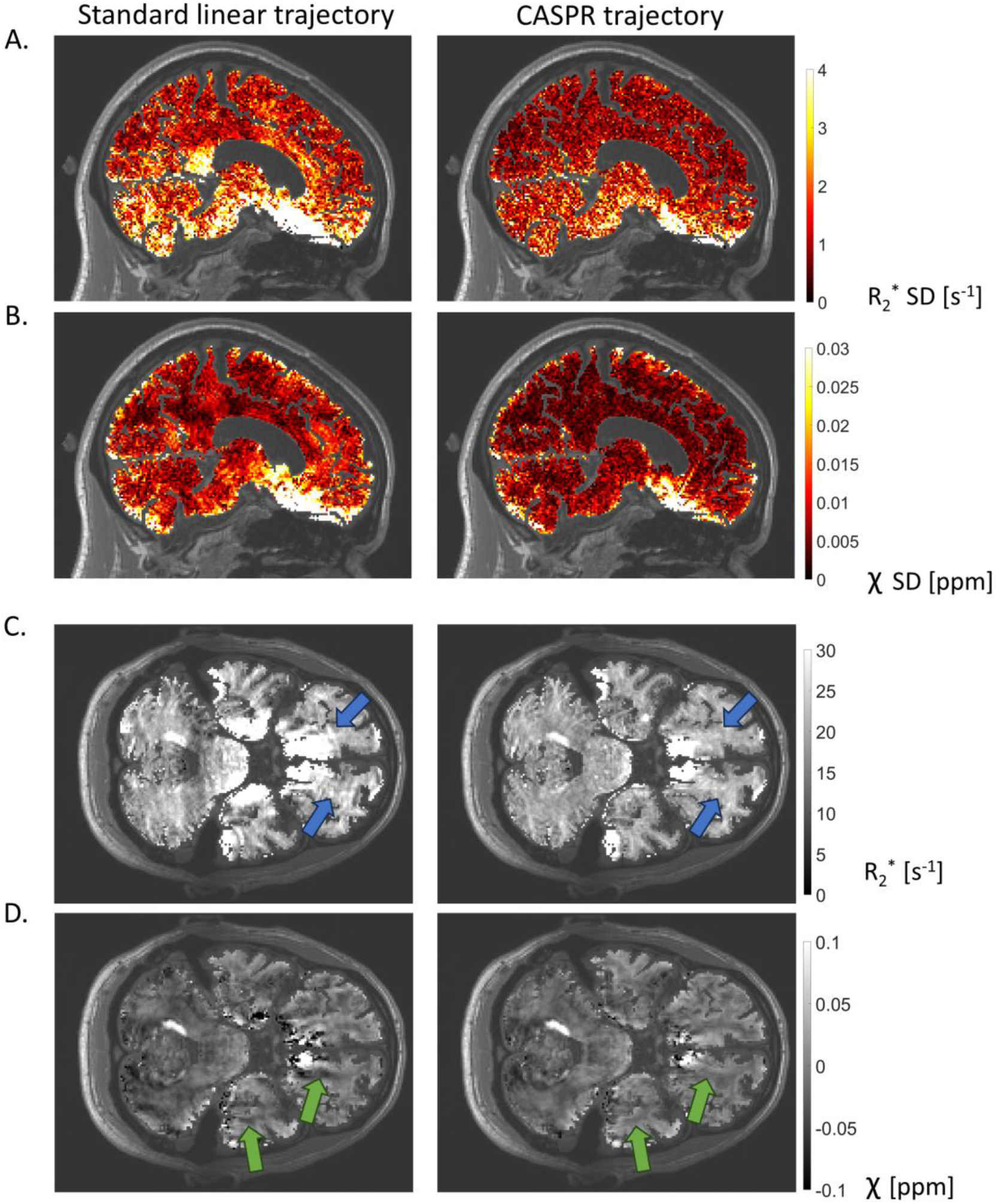
Maps of R2* and 𝜒 computed from higher resolution data acquired with the standard linear and CASPR trajectories. (A) Standard deviation (SD) of R2* across repetitions. (B) SD of 𝜒 across repetitions. (C) R2* map from one repetition. (D) 𝜒 map from one repetition. Lower SDs can be observed throughout the brain for the CASPR sampling. The blue and green arrows highlight coherent aliasing along the anterior-posterior direction, observable in the data from the standard linear trajectory but absent in the data from the CASPR trajectory.

## 4. Discussion

In this work, we proposed two new sampling strategies that aim to reduce the level of cardiac-induced noise in brain maps of R2* and magnetic susceptibility. These strategies were derived from a recent characterization of cardiac-induced noise in brain multi-echo GRE data.^41^ The first sampling strategy acquires a specific number of averages at each k-space location that was determined from the local level of cardiac-induced noise. The second sampling strategy uses cardiac triggering to synchronize data acquisition with the cardiac cycle of the participant and acquire the most sensitive region of k-space in the part of the cardiac cycle where MRI signal is most stable. Both sampling strategies primarily target the k-space centre, which contains most of the cardiac-induced noise.^41^ The ability of both proposed sampling strategies to reduce cardiac-induced noise was assessed from estimates of the transverse relaxation rate (R2*), fitting residuals (RMSE) and magnetic susceptibility (𝜒) computed from the multi-echo data.

With the standard linear trajectory, the timing of data acquisition within the cardiac period varies across repetitions, enhancing the variability of the R2* and 𝜒 estimates.^41,42^ The variability of R2* maps was 1.8-2.2s^-1^ across repetitions, higher than the variability of R2* across the cardiac cycle in the original characterization of cardiac-induced noise (0.8-1.4s^-1^).^41^ However this variability includes contributions from all noise sources, including thermal noise, which is higher here due to the higher resolution (2x2x2mm^3^ instead of 2x4x4mm^3^). Additionally, the current data displays a substantial amount of spatial aliasing along the slow phase-encoding direction (anterior-posterior in Figure 3A), leading to increased variability away from the source of cardiac-induced noise. This aliasing was not present in the original characterization of cardiac-induced noise because the k-space data was binned according to its cardiac phase before image reconstruction^41^.

The CASPR trajectory reduces the variability of R2* maps across repetitions by 26/28/22% in the brainstem/cerebellum/whole-brain. Similarly, the variability of 𝜒 maps is reduced by 19/18/16% with CASPR. These reductions are stronger than those expected from the higher number of samples required with CASPR, if thermal noise dominates signal variance. This underlines the benefit of a strategy that targets the centre of k-space, where most of the cardiac-induced noise is located. However, these numbers are lower than the fraction of the variability of R2* attributed to cardiac pulsation in the original study (∼35%).^41^ Similarly, CASPR reduces the variability of RMSE across repetitions by 17/17/13%, lower than the fraction of the variability of RMSE attributed to cardiac pulsation in the original study (∼29%).^41^ These findings are consistent with the lower impact of physiological noise in the high-resolution data presented here, although the CASPR trajectory may have allowed for the correction of other types of physiological noise such as respiration.^85^

Cardiac triggering was not effective at reducing the variability of R2*, 𝜒 and RMSE estimates across repetitions. Care was put into getting high-quality pulse oximeter signals and all post-hoc analyses indicate that cardiac triggering performed as expected. The number of cardiac triggering lines was set to 2, deemed sufficient to have a reliable and efficient synchronization (Figure 2). More cardiac triggering lines could have led to a better synchronization, at the cost of a longer scan time. Also, pulse oximeters might be too unreliable to accurately measure the phase of the cardiac cycle, resulting in a bad synchronization between the acquisition and the real cardiac phase.^87^ Moreover, even under the assumption that the synchronization was ideal, the initial assumption that cardiac-induced noise is consistent across heartbeats might not always hold: changes in cardiac-induced noise between heartbeats may lead to signal instabilities sufficient to counteract the intended benefits of cardiac triggering. Also, suspension of data acquisition may lead to discontinuities of the head position between data points acquired consecutively, resulting in enhanced aliasing.^86^

One limitation of CASPR trajectories is that they can induce stronger eddy current artefacts than their linear counterparts, due to the faster traversal of k-space along the two-phase encoding directions.^86–90^ Eddy currents can be mitigated by reducing the amplitude or slew rate of the encoding gradients, increasing the TR, or acquiring more points on each spiral arm to reduce the k-space distance between consecutive points. Moreover, scanner heating leads to a drift of the main magnetic field that can degrade the quality of data acquired with CASPR trajectories. This effect can be corrected using phase navigator data acquired throughout the scans.^91^

The proposed CASPR trajectory is a passive averaging approach that does not target specifically cardiac-induced noise and acts on other sources of physiological noise such as motion and breathing. CASPR trajectories render physiological noise spatially incoherent: physiological noise does not remain localized at its source but spreads across the 2D plane of the phase-encoding directions.^47^ Numerous methods exist to filter out incoherent aliasing artifacts, creating an opportunity for further data quality improvement or higher acceleration factors.^92–94^ Advanced reconstruction strategies such as compressed sensing^92,95^ or HD-PROST^93^ synergise well with CASPR trajectories which can be used to perform an incoherent under-sampling of the k-space data. The spatial scrambling of physiological noise with CASPR does not take place with the ‘ISME’ approach, which focuses on the temporal dimension of the multi-echo data alone.^42^ Both approaches lead to comparable reductions of the variability of R2* and 𝜒 across repetitions. One noticeable difference is that CASPR reduces the error level on the R2* estimates (RMSE) by ∼5% in inferior brain regions, while ISME increases the RMSE by ∼15%.^42^ We attribute this difference to the fact that with CASPR the k-space centre data, which contains most of the cardiac-induced noise, is acquired at multiple phases of the cardiac cycle, for all echo times of the data.

## 5. Conclusion

In this work, we evaluated two candidate strategies that aimed to reduce the level of cardiac-induced noise in brain maps R2* and magnetic susceptibility (𝜒). Both sampling strategies were motivated from a previous characterization of cardiac-induced noise in brain 3D multi-echo GRE data.^41^ The first sampling strategy relies on a CASPR trajectory to acquire a number of averages at each k-space location that depends on the local level of cardiac-induced noise. The second strategy uses cardiac triggering to synchronizes data acquisition with cardiac pulsation in real time to acquire the most sensitive area of k-space during the most stable part of the cardiac cycle.

Compared to standard linear trajectory, cardiac triggering does not efficiently reduce the variability of R2* and 𝜒 estimates across repetitions. Conversely, the CASPR trajectory reduces the variability of R2* and 𝜒 estimates across repetitions by 15-30%, for an increase in scan time of 14%. The largest improvements in variability take place in inferior brain regions such as the brainstem and cerebellum. This strategy further improves the reproducibility of R2* and 𝜒 maps by reducing the aliasing of cardiac-induced noise across the field of view, away from its original location.^91–94,96^

## Data availability statement

One of the 5D datasets used for the optimization of the sampling strategies can be found here (DOI:10.5281/zenodo.7428605). The high-resolution dataset is available online here (DOI:10.5281/zenodo.12685105).

## Funding information

This work was supported by the Swiss National Science Foundation (grant no 320030_184784 and CR00I5-235940 to AL; 32003B_182615 and CRSII5_202276 to RBvH) and the Fondation ROGER DE SPOELBERCH.

## Ethics approval statement

This study received approval from the local ethics committee and all participants gave their written informed consent prior to participation.

## 6. Supplementary material

### 6.1. Analysis of the cardiac-triggered sampling

Further analyses were performed to investigate why cardiac-triggered sampling did not reduce the variability of the R2*/𝜒 estimates across repetitions. First, an examination of the finger pulse-oximeter data was conducted, to verify that cardiac triggering was performed as expected.

#### 6.1.1. Examination of the pulse-oximeter data

The pulse-oximeter signals recorded at the participants’ finger were saved after acquisition of the MRI data. Visual inspection of each pulse-oximeter data was performed. The signal was clean and the peaks of the pulse wave seemed to have been accurately detected. Figure S1A shows a typical time-course of pulse-oximeter signal, with the corresponding periods of data acquisition and suspension. Data acquisition was stopped when reaching the cardiac triggering lines, only to be restarted when reaching detecting the next peak of the pulse wave. The peak of the pulse wave detection was conducted in real time by the MR sequence, from the detection of a local maximum in the pulse-oximeter signal with an amplitude above 2500 (the mean value of the pulse-oximeter signal). Even if simplistic, this method gives very accurate results and does not trigger on the second peak that happen during the diastole, after the dicrotic notch.

Combining the pulse-oximeter data with the k-space trajectory allows examination of the cardiac phase at the time of acquisition of each k-space data point (Figure S1B). Along the fast phase-encoding direction, the time span between consecutive points (TR=40ms) is small compared to the cardiac period and the cardiac phase varies smoothly. For a standard linear trajectory (left), the phase of the cardiac cycle shows abrupt changes along the slow phase-encoding direction. As expected with cardiac triggering (right), the phase of the cardiac cycle is 0 (peak of the pulse wave detection) at the cardiac triggering lines. As a result of the cardiac triggering lines, the phase of the cardiac cycle is highly consistent along the slow encoding direction, at the centre of k-space. At the centre of k-space, the phase of the cardiac cycle shows little variability across repetitions and participants (Figure S1C).

According to these analyses, the cardiac triggering seems to have worked as expected. The peaks of the pulse wave were accurately detected and the centre of k-space is acquired in the first quarter of the cardiac cycle, which is the least noisy part of the cardiac-cycle. The poor performance of the cardiac triggering sampling doesn’t seem to originate from an implementation problem.

**Figure S1:**
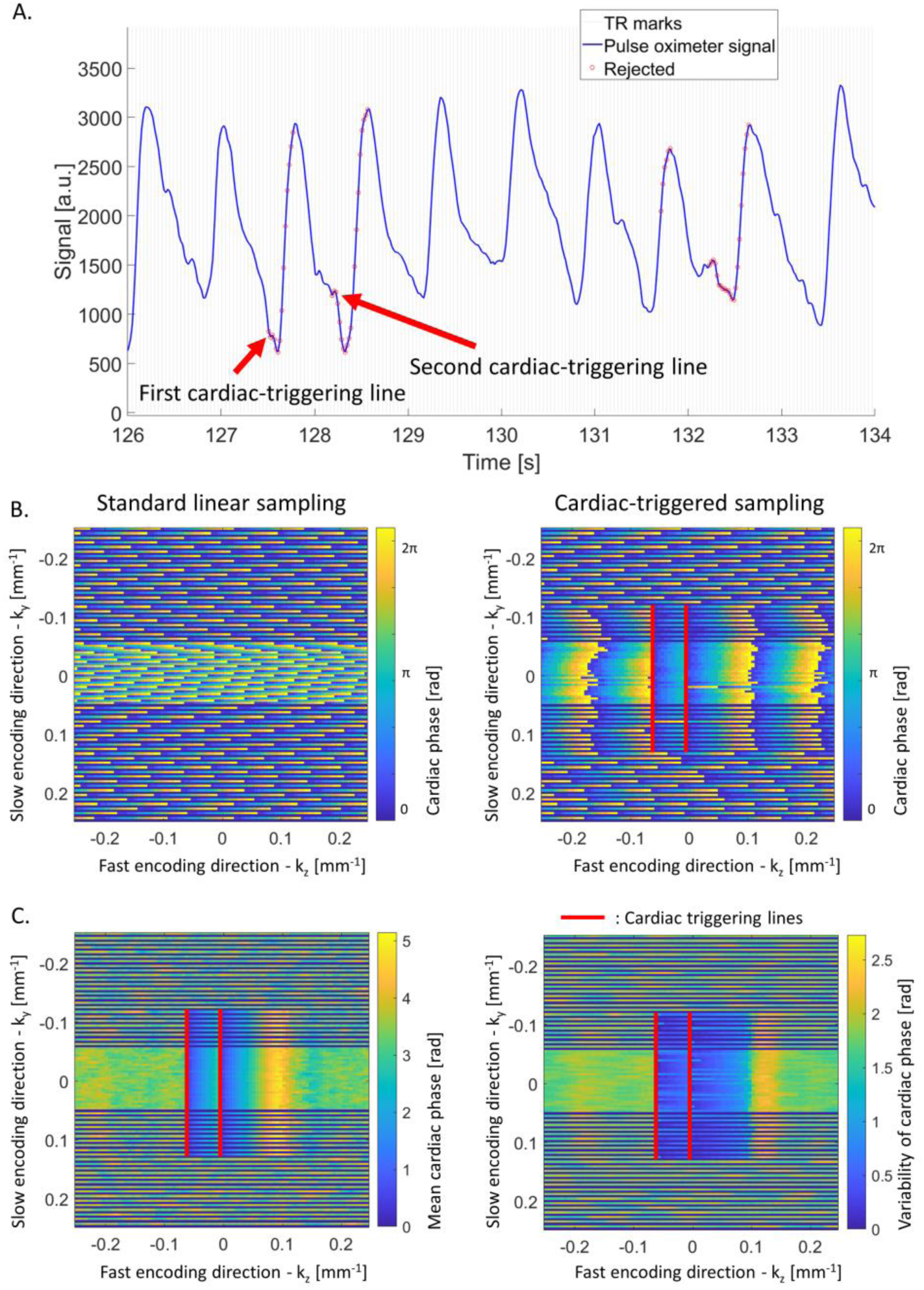
Analysis of the performance of cardiac triggering. (A) Representative example of pulse oximeter signal (blue plot). The grey vertical lines show consecutive blocks of multi-echo readouts (TR). The first and second triggering lines are indicated with arrows, and the time points where data acquisition was suspended are highlighted with red circles. (B) Example distribution of the phase of the cardiac cycle during the acquisition of the k-space data. Along the fast phase-encoding direction, the time span between consecutive points (TR=40ms) is small compared to the cardiac period and the cardiac phase varies smoothly. For a standard linear trajectory (left), the phase of the cardiac cycle shows abrupt changes along the slow phase-encoding direction. For the cardiac-triggered sampling (right), the cardiac phase is near 0 at the k-space centre and is highly consistent due to the cardiac triggering lines (red). The data were acquired with a GRAPPA acceleration factor of 2 with 24 reference lines at the k-space centre. The missing lines are shown in dark blue. (C) Mean (left) and variability (right) of the acquired cardiac phase across repetitions and participants using the cardiac-triggered sampling.

## Notes

### Competing Interest Statement

The authors have declared no competing interest.

### Summary of Updates

The title was changed and the manuscript clarity was improved.

